# Structural and functional heterogeneity in cardiac RyR signalling nanodomains revealed with quantitative single molecule mapping toolkit

**DOI:** 10.64898/2026.07.22.740006

**Authors:** Ralf Köhler, Miriam E. Hurley, Edward White, Izzy Jayasinghe

**Author notes:** These authors contributed equally (Ralf Köhler, Miriam E. Hurley).

## Abstract

The nanoscale organisation of ryanodine receptor type 2 (RyR2) channels and junctophilin-2 (JPH2) shapes cardiac Ca²⁺ release, but how this relationship is remodelled in right ventricular failure remains unclear. We developed an integrated analysis pipeline building on multiplexed DNA-PAINT data to quantify RyR2 and JPH2 abundance, stoichiometry of co-clustering, and spatial organisation within individual subsarcolemmal Ca²⁺-release nanodomains. Applied to cardiomyocytes from control rats with pulmonary hypertension-induced right ventricular failure, the approach revealed reduced co-localisation between RyR2 and JPH2 within peripheral junctions and greater variability in their co-clustering stoichiometry across the cell. A sub-variogram analysis further showed divergent remodelling of JPH2 expression patterns across subcellular length scales, indicating that disease alters both local molecular composition and cell-wide spatial heterogeneity. Experimentally derived RyR2/JPH2 maps were then used to model as two-dimensional templates for stochastic reaction-diffusion of Ca²⁺ release. Simulations of spontaneous Ca²⁺ waves from failing cells showed up to 25% wave propagation. Maps from failing cells supported faster transverse and longitudinal Ca²⁺ dependent release, consistent with the emergence, that support the likelihood of heterogeneous modulation and coupling of RyR resulting from the RyR redistribution and heterogeneous JPH2 expression in the failing cell. Together, these findings identify spatially heterogeneous RyR2-JPH2 remodelling as a potential substrate for dysregulated Ca²⁺ signalling in right ventricular failure and establish a transferable toolkit for linking molecular nanostructure to cellular function.

## Introduction

The spatial organisation of the Type-2 ryanodine receptor (RyR) channels within clusters is a major determinant of their calcium (Ca^2+^) release function. Ca^2+^ sparks may arise from either evoked or spontaneous opening of one or more RyR2 channels, recruiting adjacent RyRs through local Ca^2+^ diffusion to produce a localised Ca^2+^ release event.^1^ The fidelity, amplitude and duration of such events are therefore sensitive to the number of RyRs in a cluster, their nearest-neighbour spacing, the compactness of the array and the proximity of neighbouring clusters that may be co-recruited. Optical super-resolution and electron microscopy studies have shown that RyR clusters are heterogeneous in size and arranged into higher-order groups that may behave as functionally coupled release units.^2–6^ Subsequent electron tomography, DNA-PAINT and modelling studies challenged the idea of a simple crystalline RyR lattice, instead revealing irregular RyR arrays with gaps, variable local density and non-uniform channel adjacency.^4,7,8^ The DNA-PAINT approach was particularly pivotal in visualising individual RyR arrangement within clusters and variable co-clustering with regulatory junctional tether protein, junctophilin-2 (JPH2).

The excitability of the RyR channel is also regulated by its coupling to the wider excitation-contraction coupling machinery. In dyads and peripheral couplons, RyR clusters are positioned opposite sarcolemmal or transverse-tubular L-type Ca^2+^ channels, allowing a small trigger influx to activate a larger sarcoplasmic-reticulum Ca^2+^ release. This nanodomain architecture requires precise apposition of the sarcoplasmic reticulum and sarcolemma, maintaining a restricted junctional cleft and stable spatial relationships between RyR, L-type Ca^2+^ channels^9,10^, other Ca^2+^ handling proteins and local buffers. It is further shaped by the availability of RyR modulators, including calmodulin, FKBP12.6, calsequestrin, triadin, junctin and disease-sensitive post-translational modifications.^11,12^ Among the structural regulators, junctophilin-2 (JPH2) plays a critical role as a tether of the sarcoplasmic reticulum to the sarcolemma, supporting dyadic architecture, stabilises transverse tubules and can modulate RyR cluster organisation and signalling. ^13–15^

In cardiomyopathy and heart failure, this hierarchical regulation of RyR is disrupted at multiple levels. Failing myocytes commonly show reduced synchrony of Ca^2+^ release, slowed Ca^2+^ removal, altered sarcoplasmic-reticulum Ca^2+^ load, increased diastolic Ca^2+^ leak, and enhanced propensity for spontaneous Ca^2+^ release. These functional maladaptations occur alongside remodelling of the transverse-tubules which become sparse or disorganised, dyads which become uncoupled, RyR clusters which are redistributed or fragmented, and JPH2 expression or localisation is reduced.^6,16–18^ Super-resolution microscopy techniques have been particularly important in demonstrating that disease alters not only the abundance or phosphorylation state of RyR, but also its spatial and functional interactions with its partners.

The role of sub-cellular remodelling is less well studied in right ventricular (RV) failure caused by pulmonary artery hypertension (PAH) compared to left ventricular cardiomyopathies. Compounding the impact of the contractile deficit, the failing right ventricle is also vulnerable to arrhythmias^19^. The nanoscale architecture of Ca^2+^ release units and structural component of the cell-wide dysregulation of Ca^2+^ signalling has not been studied in depth. In the monocrotaline (MCT) induced PAH model of RV failure, right ventricular myocytes develop transverse-tubule remodelling and Ca^2+^-handling dysfunction^20,21^. Consistent with this phenotype, recent nanoscale studies have shown subcellular remodelling of the Ca²⁺-handling protein machinery and structural disruption of dyadic organisation.^21^ Molecular imaging has begun to define the RyR/JPH2 remodelling that accompanies this RV failure phenotype. In particular, the recession of JPH2 organisation from the edges of the dyadic RyR clusters was observed in MCT induced RV failure.^17^ Correlative Ca^2+^ imaging and localisation microscopy subsequently showed that spontaneous sub-sarcolemmal Ca^2+^ sparks in failing RV myocytes are associated with altered RyR cluster structure and greater heterogeneity in the relationship between local RyR number and the integral of the local Ca^2+^ spark.^22^ Together, these findings suggest that RV failure does not simply change the spatial expression of RyR uniformly, but remodels the spatial organisation, coupling and regulatory context of the Ca^2+^ release apparatus.

How RyR and JPH2 co-expression *within* the Ca^2+^ release unit are quantitatively altered within individual surface sarcolemmal Ca^2+^ -release nanodomains in RV failure has remained unresolved. In this manuscript, we test whether the resulting spatial pattern of RyR has any contribution to the altered Ca^2+^ handling and propagation of Ca^2+^ release in micron-scale regions within the myocyte. To achieve this, we present a series of emerging tools derived from single molecule localisation microscopy method known as DNA Points Accumulation for Imaging in Nanoscale Topography (DNA-PAINT), to quantify alterations in RyR/JPH co-clustering, cell-wide heterogeneity, and simulated propagation of Ca^2+^ release in the failing right ventricular myocyte.

## Results

### A quantitative Exchange-PAINT/qPAINT workflow for single RyR and JPH2 mapping

The DNA-PAINT localisation of RyR and JPH2 proteins required enzymatically isolated ventricular cardiomyocytes to be attached and fixed onto the coverslip for total internal reflection fluorescence (TIRF) microscopy (Figure 1A). The exchange-PAINT protocol for DNA-PAINT image acquisition, schematically illustrated in Figure 1B, involved immunolabelling of RyR and JPH2 proteins with antibodies carrying short, single-stranded DNA oligonucleotides with unique sequence barcodes (docking strands). In the first segment of DNA-PAINT image acquisition, the fluorescently tagged oligos (imagers), complementing the RyR-specific docking strands, were added. The stochastic and reversible binding of the imagers to their target docking strand allowed us to record single molecule fluorescence events at the RyR targets. To switch the imaging from RyR to JPH2, the RyR-specific imagers were washed out before adding the JPH2-specific Atto655 imager oligos. The localisation of events, the downstream analysis, was performed within a pipeline implemented via the open-source Python Microscopy Environment (PyME)^23^ software platform (Figure 1C). As an example, the localisations of RyR markers of a 1.2 x 1.2 µm region of interest is shown in Figure 1D. Following filtering and drift correction, the localisations are rendered as a greyscale image with an algorithm that weights local pixel intensity by the local density of markers ^24^ (Figure 1E). The sub-clusters or RyR were segmented into uniquely identifiable regions (Figure 1F) and the individual punctate densities reflecting single RyR positions were detected using a match-filter detection (Figure 1G).

**Figure 1.**
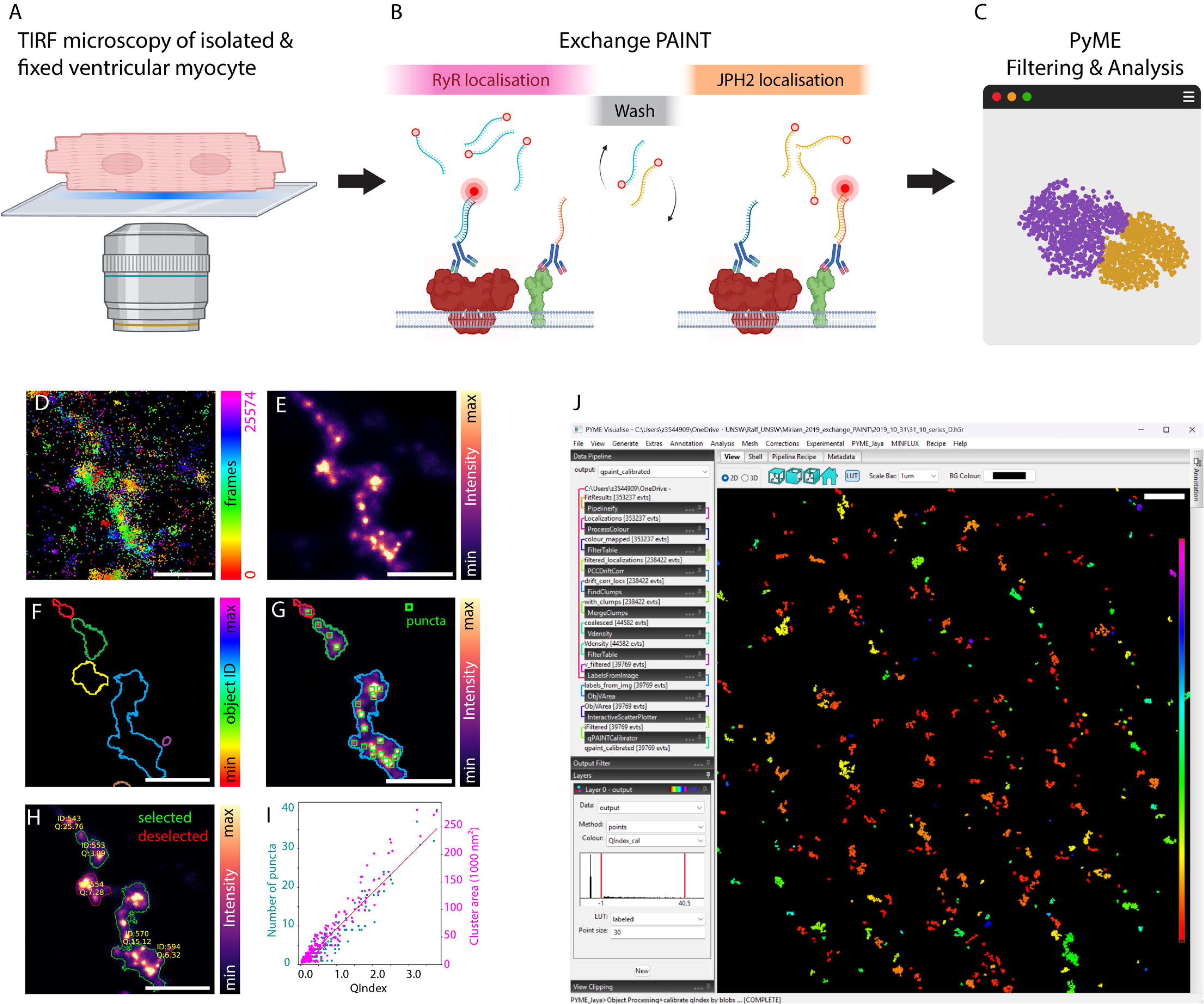
Multiplexed DNA-PAINT analysis pipeline for analysis of co-clustering properties of RyR and JPH2. **(A)** Isolated and fixed cardiomyocytes are subjected to TIRF imaging. **(B)** Under TIRF illumination, exchange-PAINT is performed by sequential RyR localisation, washing, and subsequent JPH2 localisation. **(C)** TIRF image sequences were subjected to primary event localisation, event filtering and analysis through software, Python Microscopy Environment (PyME). Shown, are exemplar visualisations of **(D)** raw single-molecule localisation events from Exchange-PAINT of RyR labelling near the cell surface, **(E)** rendered density map of the raw localisation dataset. **(F)** segmentation mask generated using an 80% signal-fraction threshold, and **(G)** matched-filter based identification of discrete RyR punct within rendered clusters; detected puncta are highlighted with green bounding boxes. **(H)** The qPAINT estimates of targets mapped on each cluster. **(I)** Direct correlation of the detected puncta within a cluster and the corresponding, Qindex-calibrated (unique) target count. **(J)** The user interface of PyMEVisualise which deploys the primary filtering and analysis pipelines on localised event data. Scale bars, 500 nm.

To move beyond visual inspection toward precise single-molecule quantification, we implemented a multiplexed analysis pipeline that consists of event filtering based on localisation parameters, drift correction, coalescence of repeated detections, segmentation, and object indexing. A detailed description of the pipeline and user-guide is provided in the Supplementary Information. Once pre-processed, the segmented regions of marker localisation in each target species were subjected to the target counting methodology, quantitative-PAINT (qPAINT) ^25^. This approach provided us an independent estimate of the number of unique RyR targets within each segmented domain (Figure 1H) which correlated directly with the number of punctate densities mapped within each RyR cluster (Figure 1I). Figure 1J shows an example of the PyME graphical user interface extended with our pipeline-specific plugins, which provides a straightforward route for applying this analysis to multiplexed RyR/JPH2 DNA-PAINT datasets with direct visualisation of cluster-specific measurements.

### Local quantification of RyR and JPH2 co-clustering

Before quantifying RyR/JPH2 co-clustering, we first examined whether the morphology of labelled junctional domains differed between conditions. Morphological analysis of RyR and JPH2 labelling patterns indicated that MCT-RV samples contained a greater proportion of elongated and irregular clusters, although individual clusters from Control-RV and MCT-RV cells spanned overlapping shape ranges (Supplementary Figure A-B). Consistent with previous studies, RyR clusters in MCT-RV cells were typically smaller^17,26^.

To capture the differences in RyR and JPH2 co-expression *within* the clusters, we used the qPAINT analysis pipeline to count of the two species of targets within the bounds of the segmented clusters. Representative images of RyR/JPH2 clusters from Control-RV and MCT-RV cells are shown in Figure 2A and 2B respectively. Analysis of the cluster-specific ratio between the qPAINT counts of RyR and JPH2 targets showed a median decrease of 14% between Control-RV and MCT-RV (Figure 2C). A noted feature of this analysis was also the higher variance in this ratio in MCT-RV clusters, suggesting that the reduction of their co-clustering stoichiometry is not observed homogeneously across the sub-sarcolemmal regions imaged across the diseased cells. In the scattergrams correlating the RyR QIndex values with the cluster-matched JPH QIndex (Figure 2D), we also observe a reduced correlation coefficient in MCT-RV samples compared to Control-RV (R^2^ of 0.528 compared to 0.383).

**Figure 2.**
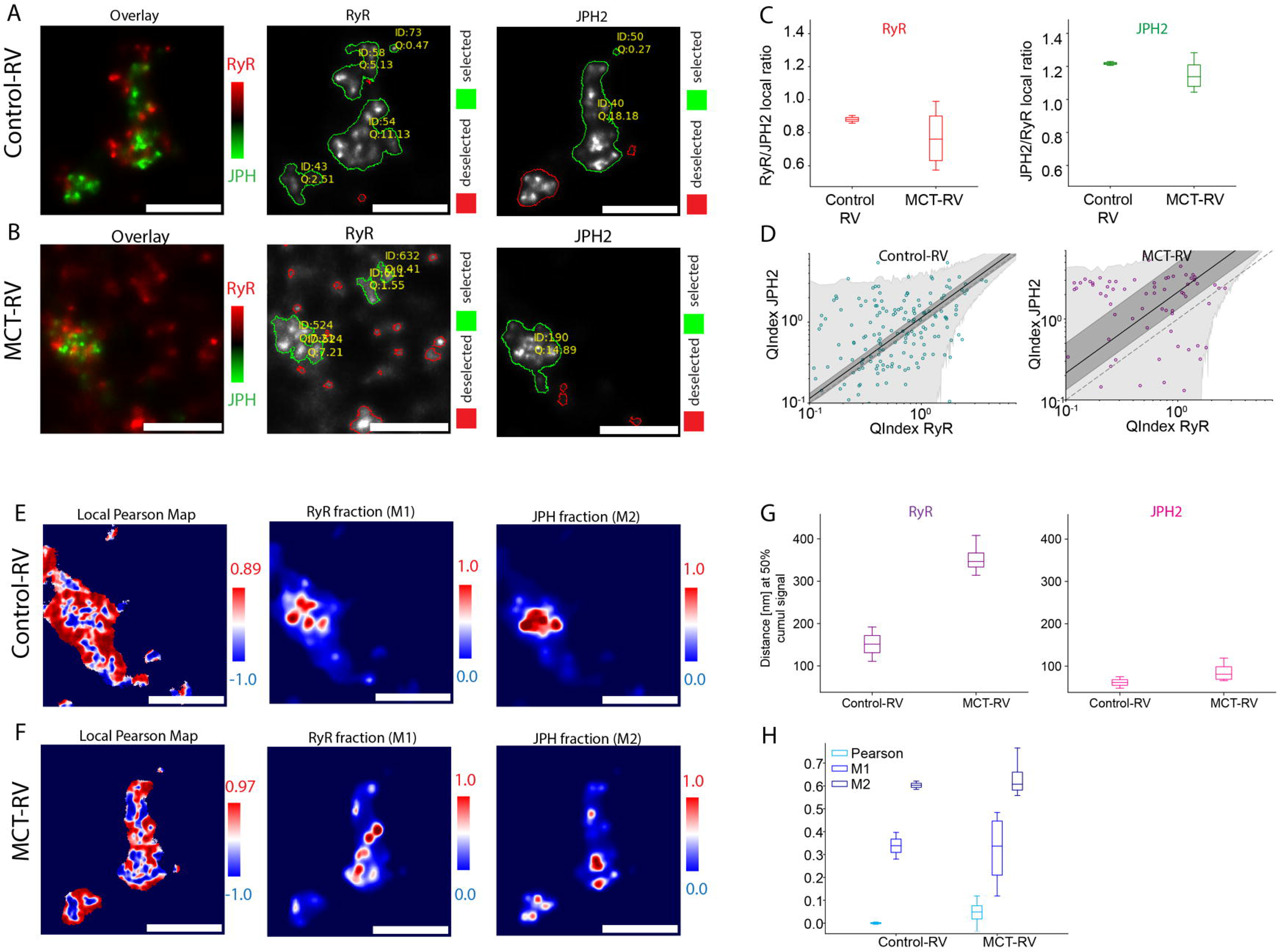
Analysis of molecular-scale interactions between RyR and JPH within junctions. Shown, are representative two-colour overlays, RyR (red), and JPH (green) nanodomains from **(A)** Control-RV and **(B)** MCT RV samples. In the clusters featuring DNA-PAINT localisations from both targets, the cluster identifier and local QIndex units are annotated. **(C)** Cluster-wise stoichiometric ratios derived from qPAINT-calibrated site counts. Left: JPH2:RyR (number of JPH2 targets per RyR2 target). Right: reciprocal RyR2:JPH2 ratios. Boxplots summarise quartiles for images analysed in Control-RV and MCT-RV datasets. **(D)** Object-level QIndex comparison for paired RyR2 and JPH2 clusters for Control-RV (right) and MCT-RV (left). QIndex values for overlapping clusters were plotted against each other within each image; each point represents one paired object. The solid line shows the best-fitting linear relation, with the dark grey band indicating the 95% confidence interval for the fit and the light grey band the 95% prediction interval. The dashed line denotes the 1:1 relationship. Local Pearson correlation map (left), calculated with a 35 x 35 nm (7 x 7 pixels) moving window between RyR2 and JPH2 signals, and local Manders M1 and M2 coefficient maps are shown for exemplar **(E)** Control-RV and **(F)** MCT-RV images. (G) Box and whisker plots illustrate the quartiles of the measurements of the distance from the JPH densities within which 50% of the cumulus of RyR signal was localised (left) and vice versa (right) in Control-RV and MCT-RV. (H). A summary box plot comparing the quartiles of the measurements of Pearson’s correlation co-efficient, Manders M1, and M2 coefficients. Scale bars: 500 nm.

Adjacent to the qPAINT ratio analysis, Person’s correlation and Manders co-localisation analyses allowed us to quantify the spatial shifts in the JPH2 organisation relative to RyR. To visually represent these measurements across the super-resolution images, Pearson’s correlation co-efficient and Manders co-efficients (M1: representing the ratio of RyR co-localising with JPH and M2: vice versa) were computed within a 35x35 nm (7x7 pixels) moving window for representative Control-RV and MCT-RV images (Figure 2E & 2F respectively). Whilst the median Manders co-efficients show negligible change, the variance between measurements in MCT-RV was 2.3-fold higher compared to Control-RV (see summary box plot in Figure 2H). A Euclidean Distance Transform was used to quantify the distance at which 50% of the co-label was observed from a given protein species. This analysis showed over a 2-fold increase in the distance within which JPH2 events were found from the nearest RyR event density in MCT-RV, compared to Control-RV. The reciprocal analysis for RyR events (in relation to JPH2) showed no statistical change between the two conditions.

### Analysis of scale-specific heterogeneity of RyR and JPH2 expression

To determine whether the remodelling observed at the level of individual clusters extended to larger length scales, we implemented an analysis based on a previously developed protocol to identify spatial heterogeneities in protein expression density maps.^27^ This analysis generates a heatmap, known as a Gaussian Random Field (GRF), revealing the length-scale specific variability in the protein expression through the region of interest. In the shown GRFs, RyR and JPH2 (Figure 3A and 3B respectively) images were respectively downscaled to a series of local scale factors ranging from 100 to 1000 nm that reflect the length-scales λ. Similar to any greyscale fluorescence image, the high intensity pixels in the GRF reflect regions of high density of expression. High contrast within GRF visualisation reflects high degree of spatial heterogeneity in that specific length-scale.

**Figure 3.**
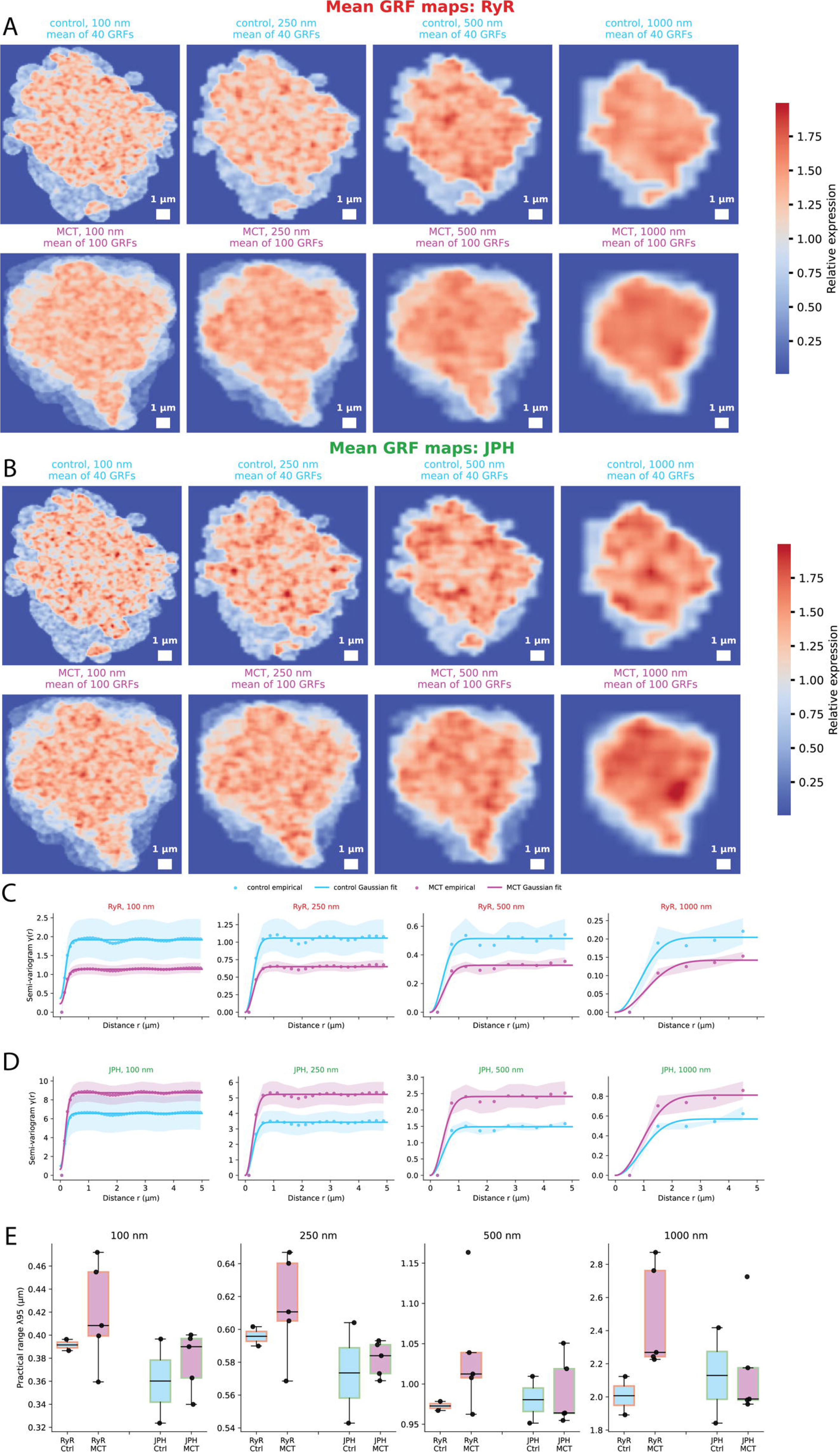
Analysis of scale-specific heterogeneity of the RyR and JPH2 organisation. Rendered RyR2 and JPH2 density maps were binned at 100, 250, 500, and 1000 nm and analysed using masked FFT-based semi-variograms. Mean Gaussian random field maps generated from fitted variogram parameters for **(A)** RyR and **(B)** JPH2, respectively. These synthetic relative-expression fields visualize the spatial scale and amplitude of heterogeneity captured by the model and are not experimental images. Shown, are average empirical semi-variograms for Control and MCT images of **(C)** RyR and **(D)** JPH. The fitted sill reflects the plateau variance of the density field, whereas λ95 denotes the practical correlation range at which the fitted variogram reaches 95% of the sill. **(E)** λ95 comparison across channels, conditions, and binning scales.

Both the contrast and the local intensity values of the GRF in RyR image data were comparable between Control-RV and MCT-RV image data across the different length scales (Figure 3A). By comparison, JPH2-derived maps showed more prominent high-density domains, suggesting divergent changes in the spatial heterogeneity JPH2 expression (Figure 3B), particularly across λ approaching 1000 nm.

To quantify these observations a series of plots, called ‘empirical semi-variograms’ were calculated to analyse the overall contrast of the GRFs (reflected by the variance of the downsampled images). For a given length-scale of interest λ, the semivariance as a function of the distance (*r*) were fitted with a Gaussian variogram model. See the detailed methodology in Supplementary Methods. Note that characteristically these plots reach a plateau as spatial correlation between points decrease with increasing distance. The practical correlation range, is defined as the distance at which the fitted semivariance reaches 95% of its plateau value. These plots revealed that for all of the considered length scales, the analysis returned lower semivariogram values for the RyR signal in MCT-RV image data compared to that in Control-RV (Figure 3C). The converse of this trend was observed for the equivalent analysis of JPH2 (Figure 3D). Whilst values for RyR and JPH2 image data generally increased with the higher length-scales, there were no statistically significant differences observed between Control-RV and MCT-RV groups of datasets.

### Simulation of spontaneous Ca^2+^ waves on experimentally derived RyR/JPH2 maps

We next sought to test weather changes in RyR cluster structure and increasingly more heterogeneous JPH2 expression distribution observed in MCT-RV cells were sufficient to alter the propagation of spontaneous Ca^2+^ waves. For this analysis, experimentally derived RyR2/JPH2 maps were used as the structural template for a stochastic reaction-diffusion simulation implemented within the PyME pipeline.

For this analysis implemented natively through the PyME pipeline, we selected two-dimensional (2D) DNA-PAINT images of sub-sarcolemmal RyR and JPH maps from exemplar Control-RV and MCT-RV cells (Figure 4A-i and 4B-i). In visual comparison, we observed the more fragmented nature of the RyR organisation and the lower density of JPH2 expression in MCT-RV images (seen in magnified views, Figure 4A-iii and 4B-iii). This data was converted to a series of white pixels (each reflecting the 2D position of detected RyR puncta), and cluster perimeters obtained by binarising the JPH2 image. The pixel values of the cluster area encoded the QIndex values of the unique JPH2 count estimated within the local cluster (Figure 4A-ii and 4B-ii, respective magnified views shown in panels iv).

**Figure 4.**
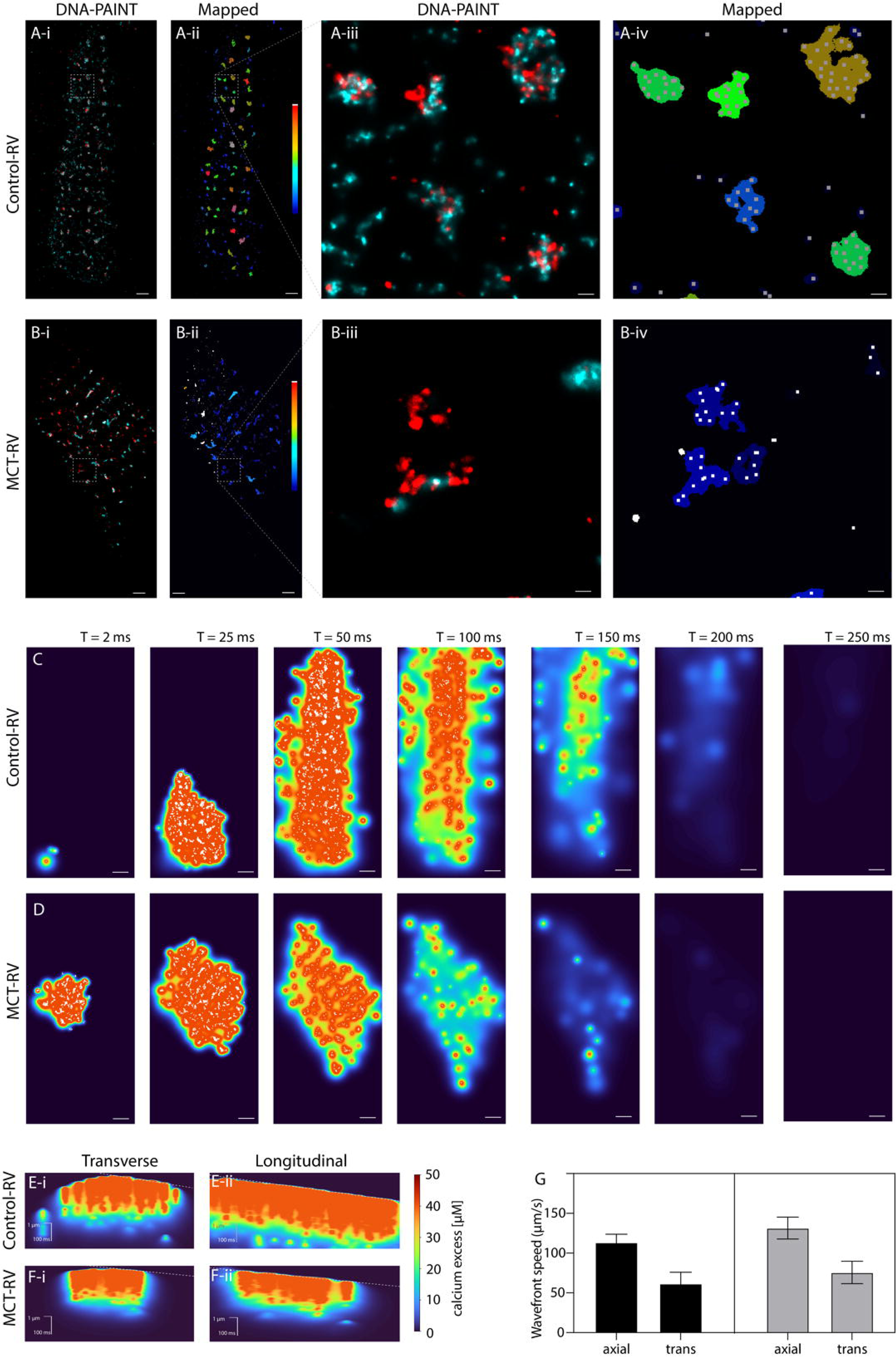
Simulation of spontaneous and propagating Ca^2+^ release on sub-sarcolemmal DNA-PAINT maps of RyR. Shown, are two exemplar exchange-PAINT images of RyR (red) and JPH2 (cyan) in **(A-i)** Control-RV and **(B-i)** MCT-RV. From DNA-PAINT images, maps of single RyR puncta (represented by white pixels) and binarised JPH2 clusters encoding the local qIndex(JPH) are constructed for **(A-ii)** Control-RV and **(B-ii)** MCT-RV respectively. Magnified views of the respective regions of interest (dashed boxes) in both the rendered images and encoded maps are shown in panels **iii** and **iv**. (**C & D**) Time interval series of the Ca^2+^ waves simulated from the Control-RV and MCT-RV maps shown above. White pixels indicate the momentarily open RyRs. Linear kymographs along the (**i**) transverse and (**ii**) longitudinal axes of the **(E)** Control-RV and **(F)** MCT-RV simulations. Dashed lines indicate the approximated speed of the wave front in the transverse and longitudinal axes. (G) Bar plots comparing the Ca^2+^ wave speed in Control-RV and MCT-RV simulations in both transverse and longitudinal axes. Scale bars. A-i&ii, B-i&ii, C&D and horizontal bars of E&F: 1 µm; A-iii&iv, B-iii&iv: 100 nm. Vertical bar in E&F: 100 ms.

The simulation featured a stochastic discrete-state model of RyR gating with Markov-like [Ca^2+^]_i_-dependent transitions (see details in Supplementary Methods section 1.2). At each timestep, closed RyRs opened according to a Monte Carlo probability determined by the local calcium concentration and JPH2-dependent sensitivity scaling. This stochastic channel model was coupled to a 2D reaction-diffusion simulation of cytosolic calcium spread which was triggered at a random location seeded within the lower left corner of the cell area within the DNA-PAINT image. Figure 4C & 4D illustrate the C^2+^ wave simulations for the Control-RV and MCT-RV datasets at chosen time-points, along with the momentary RyR opening patterns (white + marks). Kymographs taken along the axial and transverse axes of the cell allowed us to visualise the spatiotemporal profile of the wave spread (Figure 4E & 4F respectively). Regression fits to the wavefronts of these kymographs provided estimates of the axial and transverse wave speeds, collated and compared across Control-RV and MCT-RV datasets (Figure 4G). Whilst the transverse speed of wave spread was 74-84% faster than the axial spread, we also observe a 23% and 16% increase in these respective speeds in MCT-RV compared to Control-RV. The kymographs further indicated that wavefront propagation was spatially heterogeneous, with local differences in slope consistent with non-uniform recruitment of release sites during wave spread.

## Discussion

This study introduces an integrated quantitative single-molecule mapping and image-based modelling framework for examining the organisation of cardiac Ca^2+^ release nanodomains in right ventricular failure. By combining Exchange-PAINT, qPAINT molecular counting, cluster-resolved spatial analysis, variogram-based characterisation of subcellular heterogeneity, and stochastic Ca^2+^ release simulations, we examined RyR and JPH2 organisation at several spatial and functional levels. Three principal findings emerge. First, qPAINT provided a practical means of estimating RyR2 and JPH2 abundance within individual junctional domains, extending the analysis beyond cluster morphology and fluorescence intensity. MCT-induced RV failure was associated with a less consistent relationship between RyR and JPH2 abundance and nanoscale apposition, with disease-related changes expressed most prominently as increased variability between local domains. Further, when experimentally derived molecular maps were used as the structural basis of a stochastic reaction-diffusion model, the MCT-RV configuration supported faster simulated Ca^2+^ propagation than the Control-RV configuration. Together, these findings suggest that RV failure remodels not only the amount or arrangement of Ca^2+^ handling proteins, but also the local molecular context in which RyR channels are activated.

A central methodological contribution of the study is the implementation of spatial molecular counting within an open-source single-molecule analysis environment. Conventional super-resolution measurements of RyR organisation have commonly relied on cluster area, event density, puncta number or nearest-neighbour spacing. These measures are informative but can be affected by localisation oversampling, variation in labelling density and differences in image rendering. qPAINT instead uses the predictable dark-time kinetics of DNA-PAINT binding to estimate the number of accessible docking sites within a region. The relationship observed here between qPAINT-derived target estimates and independently identified RyR puncta supports the use of the approach for comparing RyR-containing nanodomains.

Embedding the workflow within PyME also provides a route towards reproducible analysis of multiplexed localisation microscopy datasets. Event filtering, temporal coalescence, density-based background removal, segmentation, qPAINT analysis and cluster-level measurements can be performed within a common data framework while retaining links between measurements and the underlying localisation events. This is valuable for cardiac Ca^2+^ nanodomains, in which apparently similar rendered structures may contain different numbers or combinations of molecular targets. Because DNA-PAINT kinetics are affected by imager concentration, temperature, sequence and acquisition conditions, calibration remains experiment-specific. The principal strength of the present approach is therefore not an assumption of universal QIndex values, but the ability to make calibrated, internally consistent comparisons between molecular species and biological conditions acquired under controlled conditions.

The cluster-level analyses indicate that RyR-JPH2 remodelling in MCT-RV is heterogeneous rather than uniform. The average stoichiometric shift between conditions was modest, but the dispersion of the cluster-specific ratios increased in MCT-RV cells. In parallel, the co-location between paired RyR and JPH2 QIndices was weaker in the diseased samples. These findings suggest that the abundance of the two proteins becomes less tightly coordinated between individual Ca^2+^ release domains. Thus, a population-average measure of RyR or JPH2 expression may underestimate the extent of local junctional remodelling. Some domains may retain an approximately control-like molecular composition, whereas others may become relatively depleted of JPH2 or display altered RyR:JPH2 stoichiometry.

This interpretation is supported by the distance-transform analysis. The increased distance required to encompass 50% of the JPH2 signal relative to RyR indicates a broader displacement or redistribution of JPH2 around RyR-containing domains in MCT-RV. By contrast, the reciprocal distance measurement showed little change. Although the directionality of these measurements depends partly on the size, density and continuity of the two labelled structures, the asymmetric result is consistent with JPH2 receding from, or becoming less concentrated around, comparatively persistent RyR domains. This interpretation agrees with previous observations of molecular-scale dyadic fraying in MCT-induced RV failure. It is also compatible with the established role of JPH2 in maintaining junctional membrane complexes, stabilising transverse-tubular organisation and regulating the functional environment of RyR channels.

The limited changes in global Pearson and median Manders coefficients illustrate why multiple spatial measurements are required. Global colocalisation coefficients compress structurally complex images into single summary values and may be relatively insensitive to a redistribution that affects only a subset of junctions. In the present data, the increased variance of the local measurements and the altered distance relationships were more prominent than changes in their central tendency. The disease phenotype may therefore be characterised less by complete separation of RyR and JPH2 than by an increase in the frequency and severity of locally mismatched domains. Such heterogeneity could be functionally important because Ca^2+^ release is determined locally: a small number of structurally or molecularly unstable junctions may contribute disproportionately to spontaneous release even when the average organisation across a cell appears only moderately altered.

The variogram analysis extends this conclusion from individual clusters to larger subcellular length scales. RyR2 and JPH2 displayed divergent changes in semivariance between Control-RV and MCT-RV images. RyR2 maps generally showed lower semivariogram amplitudes in MCT-RV, whereas JPH2 maps showed the opposite tendency. However, the fitted practical correlation ranges did not differ significantly between groups. This combination suggests that RV failure primarily altered the magnitude of spatial density fluctuations represented within the analysed length scales, rather than producing a clearly resolved shift in the characteristic size of the correlated domains. In other words, the extent to which local protein density deviated from the cell-wide mean changed more evidently than the distance over which those deviations remained spatially correlated.

The divergent behaviour of the two channels further suggests that JPH2 may be a particularly labile component of the failing junction. RyR clusters can fragment, redistribute or change in density during disease, but they remain anchored within a broader sarcoplasmic-reticulum network. JPH2 organisation is additionally dependent on the integrity of membrane contacts and the transverse-tubule system. It may therefore undergo spatial redistribution over both nanometre and micrometre scales as junctional membranes remodel. The qPAINT, distance-transform and variogram measurements provide complementary views of this process: altered stoichiometry within individual clusters, altered nanoscale apposition between the proteins and altered larger-scale fluctuations in JPH2 density.

The functional significance of this heterogeneous remodelling was explored using experimentally derived RyR2 and JPH2 maps as the spatial substrate for Ca^2+^ release simulations. Within the model, the MCT-RV map supported faster propagation in both the transverse and longitudinal directions. Propagation was also faster transversely than longitudinally in both conditions, indicating that the anisotropic spatial organisation of release sites remained an important determinant of the simulated wavefront. These observations show that realistic molecular maps can produce condition-dependent emergent behaviour even when the underlying gating and diffusion rules are otherwise held constant.

At first sight, faster propagation from the more fragmented MCT-RV architecture may appear counterintuitive. Fragmentation and loss of local RyR density would be expected to reduce the probability of coordinated recruitment within some clusters and could decrease the amplitude or fidelity of an individual Ca^2+^ spark. However, local release synchrony and long-range wave propagation are not equivalent properties. A heterogeneous network containing sensitised RyRs, variable JPH2 association and irregularly spaced release sites may support saltatory recruitment through a subset of highly excitable domains. In such a system, some junctions may function poorly during triggered excitation-contraction coupling, while others may provide preferential routes for spontaneous propagation. This offers a possible framework for reconciling impaired synchronous Ca^2+^ release in failing myocytes with an increased susceptibility to spontaneous events and arrhythmogenic Ca^2+^ waves.

The simulation should nevertheless be interpreted as a mechanistic test of the consequences of the mapped architecture rather than as a complete physiological reconstruction. JPH2-dependent modulation of RyR sensitivity was imposed as a model relationship based on the proposed stabilising influence of JPH2; it was not measured directly for each junction. Therefore, the faster propagation in the MCT-RV simulation reflects the combined effects of the experimentally observed molecular maps and the specified relationship between local JPH2 abundance and RyR excitability. The result supports the hypothesis that heterogeneous RyR2-JPH2 remodelling can affect propagation, however its manifestation in the living myocyte in combination with the myriad of other changes in the cell’s Ca^2+^ handling proteins and homeostasis remains to be tested.

Several other limitations should be considered. Exchange-PAINT imaging was performed close to the cell surface under TIRF or highly inclined illumination and therefore primarily sampled peripheral couplings rather than the full three-dimensional dyadic network. The molecular counts represent antibody-accessible docking strands and are influenced by epitope accessibility, conjugation efficiency and steric effects. Although kinetic filtering, sample-specific calibration and comparison with puncta counts reduce these uncertainties, qPAINT estimates should be interpreted as calibrated target estimates rather than exact counts of all endogenous protein molecules. Cluster segmentation may also merge closely apposed structures or divide irregular domains, affecting the assignment of RyR2 and JPH2 events to individual objects.

The MCT model reproduces important features of pulmonary hypertension and RV failure but does not encompass the full diversity of human right-sided heart disease. The present experiments were also performed in male animals, and sex-dependent differences in RV remodelling were not addressed. Statistical inference should remain based on the biological sampling unit rather than the very large number of clusters generated by each image. Further work using larger cohorts, additional models and human tissue will be required to determine which aspects of the observed RyR2-JPH2 phenotype are conserved across different causes and stages of RV failure.

It should be noted that the Ca^2+^ wave model is not a mechanistically complete simulation of the working cell. It omits several determinants of release, including three-dimensional junctional geometry, local sarcoplasmic-reticulum Ca^2+^ content and depletion, spatial variation in SR Ca^2+^ ATPase and Na^+^/Ca^2+^ exchanger activity, L-type Ca^2+^ channel organisation, membrane voltage and the effects of other RyR-associated proteins and post-translational modifications. Where simulations were generated from exemplar molecular maps, repeated stochastic iterations quantify uncertainty in channel recruitment on those maps but do not substitute for replication across independently imaged cells. Direct experimental measurements of Ca2+ wave initiation and propagation in cells subsequently mapped by multiplexed DNA-PAINT would provide an important validation of the model predictions.

Despite these limitations, the study provides a framework for linking molecular-scale protein organisation to emergent Ca^2+^ signalling behaviour. The findings suggest that RV failure is associated with a breakdown in the locally coordinated organisation of RyR and JPH2, expressed as increased variation in molecular stoichiometry, altered nanoscale apposition and divergent spatial heterogeneity. The simulations further show that these experimentally observed architectures are sufficient, within a defined biophysical model, to alter the spread of spontaneous Ca^2+^ release. More broadly, the integration of molecular counting, spatial statistics and image-based simulation offers a transferable approach for studying how heterogeneous remodelling of signalling nanodomains contributes to cellular dysfunction.

## Methods and materials

### Exchange-PAINT labelling and imaging

Experiments were performed according to the UK Animals (Scientific Procedures) Act of 1986 and with UK Home Office approval (license number 70/8399) and approval of the University of Leeds Ethics Committee. As a model of RV failure, adult male Wistar Crl rats aged ∼5 weeks were given an intraperitoneal injection of crotaline (Merck, NJ) to induce PAH. Age and sex-matched controls were given an equivalent bolus of saline. At 21-28 days, the MCT treated animals were monitored for signs of RV failure and euthanized along with age-matched Ctrl animals. Hearts were dissected acutely and right ventricular myocytes enzymatically isolated following Langendorff perfusion. See the Supplementary Information section for details of the cell isolation procedure.

### DNA-PAINT

#### Microscope set up and image acquisition

Experiments were performed on a modified TE2000 TIRF microscope (Nikon, Japan) equipped with a 800 mW 642 nm diode laser (Viasho, China) focused onto a 15 x 15 µm field of illumination using a 1.49 NA 60x TIRF objective lens (Nikon). A Chroma T685lpxr 1 mm dichroic mirror used to split the emission light from the excitation path. Emission filter ET720/60m (Chroma) was placed in the emission light path, in front of the Zyla 5.5 USB scientific CMOS camera (sCMOS; Andor, Belfast) used to record the image data. The microscope and camera were controlled using a Lenovo Thinktank workstation using an Intel i7 quad-core processor, 32 Gb of DDR3 memory and a 2 Tb solid-state drive running the open-sources Python Microscopy Environment (PyME) software (available open-source via www.python-microscopy.org). The ‘PyMEAquire’ image acquisition interface included a live display of the camera’s acquisition at a customizable frame-rate. This was particularly critical for acquiring reference images of the sample between steps, for orientating the sample and for positioning the cells’ region of interest within the illumination TIRF field.

Primary antibodies against RyR2 and JPH2 were conjugated to single-stranded DNA docking strands using maleimide-thiol chemistry following Jayasinghe et al. and Hurley. Conjugates were purified by size-exclusion chromatography and stored at −20 °C. Fixed cells were incubated overnight at 4 °C with each DNA-antibody conjugate (1:100-1:200) in PBS + 3% BSA, washed extensively in high-salt buffer (PBS with 500 mM NaCl), and mounted in an oxygen-scavenging imaging buffer containing Trolox, pyranose oxidase, and catalase.

Dual-colour Exchange-PAINT imaging was performed on a modified NIKON TE2000 TIRF microscope (60×, 1.49 NA oil objective; 647 nm excitation) under oblique or total internal reflection illumination. Complementary Atto655-labelled imager strands were added sequentially (typically 1.2 nM) to image RyR2 and JPH2 in separate acquisition rounds (20000-30000 frames per channel, 100 ms exposure). Between rounds, samples were washed in Buffer C to remove imagers. Image sequences and localisation metadata were acquired in PYME’s native .h5r format.

### Event localisation and primary processing

Single-molecule localisations were obtained in PYME by fitting 2-D Gaussians to detected spots. For each localisation, we extracted x and y positions, localisation error, fitted amplitude, frame, and fitted sigma. A first-stage quality filter excluded events with large localisation error, low amplitude, or extreme spot width, corresponding to poorly fit or dim molecules, followed by drift-correction (Figure 1D).

To reduce oversampling of individual docking strands, temporally adjacent events within a radius equal to the mean localisation error and within a 300 ms time window were coalesced into a single “binding event”. This produced a list of temporally distinct binding events per docking strand, reducing artificial inflation of event counts arising from rapid rebinding

Lateral drift was corrected using a piecewise linear algorithm based on fit parameters extracted from the acquisition metadata; when such parameters were unavailable, we instead applied phase cross-correlation registration (Van Der Walt et al., 2014) between time-binned image projections.

Residual low-density background events were removed using Voronoi-based density filtering. First-rank Voronoi polygons were constructed for all events, and points residing in regions with local density below 10% of the global mean were discarded, following Levet et al. (Levet et al., 2015). This adaptively suppresses spatially isolated events while preserving densely packed domains).

### Click or tap here to enter text

#### Rendering, segmentation, and cluster-level feature extraction

Filtered events were rendered into super-resolution images (Figure 1E) using a jittered-triangulation algorithm in PYME (5 nm pixel size, 100 jitter iterations). The jitter amplitude ranged to the local nearest-neighbour distance to avoid over-smoothing dense regions while still suppressing sampling artefacts in sparse areas (Baddeley et al., 2010).

Rendered maps were segmented in PYME using one of the implemented intensity-based thresholding options, including cumulative signal-fraction, isodata, or log-isodata thresholding. For each rendered image, candidate masks were inspected visually against the underlying localisation density map, and the thresholding option was selected to separate adjacent high-density clusters while preserving clearly continuous signal domains. The selected binary mask was then used for connected-component labelling, with components larger than 500 nm² retained as individual clusters (Figure 1E). The segmentation method and parameter values used for each image were recorded to ensure reproducibility.

For each cluster, we computed basic geometric properties using custom Python scripts and functions from the scikit-image library (Van Der Walt et al., 2014), including area, centroid position, and the lengths of the major and minor axes of the best-fitting ellipse. Aspect ratio was defined as the ratio of major to minor axis length, and circularity as 4π×area/perimeter^2^. These metrics were used to compare cluster morphology between conditions (Figure 2E, 2F).

Rendered images were generated for each channel in PYME, and the BlobFinder module was applied to detect local intensity maxima corresponding to individual binding sites (“puncta”, Figure 1G). Briefly, BlobFinder identifies suprathreshold peaks in the density map and returns their centroid positions, enforcing a minimum separation so that closely spaced peaks are not counted multiple times. For each cluster, we then computed pairwise Euclidean distances between all puncta centroids and extracted the first- and fourth-order nearest-neighbour distances (NN1 and NN4) for each punctum. These nearest-neighbour distances were summarised per cluster and used as simple measures of local expression patterns and regularity (Supplementary Figure C-i and C-ii).

### qPAINT index calculation, calibration, and kinetic selection

qPAINT indices were computed from the distribution of dark times between binding events. For each cluster, the cumulative distribution of dark times was fitted with a mono-exponential decay, and the inverse of the characteristic dark time (τ_dark⁻¹) was taken as the qIndex. All datasets were acquired under identical imager concentration and imaging conditions to ensure comparability.

To identify clusters that comply with the qPAINT model, we plotted per-cluster qIndex against projected area for each object-related event (Figure 1I). Clusters following the expected linear relationship between qIndex and area, consistent with an approximately constant docking-site density across clusters, were then selected using a new interactive scatter-selection module (“InteractiveScatterPlotter”) that we implemented in the PYME recipe framework. This tool allows the user to lasso-select points in qIndex-area space (or any other feature of interest) and writes the selection back into the underlying localisation table as a boolean identifier, so that all subsequent stoichiometric analyses are restricted to qPAINT-consistent clusters. This step removes objects dominated by kinetic outliers, merged structures, or poorly sampled domains. The same interface can also support qIndex calibration by linking qIndex values to independently detected puncta or blob counts.

### Stoichiometric ratios and colocalisation analysis

For each segmented cluster, qPAINT-derived binding-site numbers were computed as qIndex multiplied by the calibration factor for that imaging session. Stoichiometric ratios were then expressed as JPH2:RyR2 (number of JPH2 sites per RyR2 site) and the reciprocal RyR2:JPH2 ratio (Figure 2C). The standard deviation of ratios across clusters within each cell was used as a within-cell heterogeneity measure (Supplementary Figure D-i and D-ii).

Spatial overlap was quantified using two complementary approaches. First, Euclidean Distance Transform (EDT) maps were computed for binary masks of each channel. For a given radius r, the fraction of RyR2 (or JPH2) signal lying within distance r of the other channel was measured, and the distance at which this fraction reached 50% (EDT-50) was taken as a summary measure of nanoscale apposition (Figure 2G). Second, Pearson correlation coefficients and Manders overlap coefficients were computed from rendered images of each field of view, providing global colocalisation metrics (Figure 2H). Local sliding-window Pearson and Manders maps were additionally calculated for visualisation of within-cluster variation in colocalisation (Figure 2E-F).

### Variogram analysis of spatial heterogeneity

Spatial heterogeneity in rendered RyR2 and JPH2 density maps was quantified using a variogram-based approach adapted for cardiomyocyte nanostructure analysis by Holmes et al ^27^. Rendered RyR2 and JPH2 images were analysed separately. A shared biological mask was generated from the combined density maps to exclude empty background regions, after which images were binned to multiple spatial samplings, 100, 250, 500 and 1,000 nm/pixel. Within the mask, density maps were normalised to a mean intensity of one to generate relative-density fields.

Spatial autocorrelation was estimated using a masked FFT-based autocovariance approach, allowing correction for finite image size and irregular mask geometry. Radially averaged autocorrelation profiles were converted into empirical semi-variograms and fitted with a Gaussian variogram model. The fitted parameters were used to estimate the nugget, partial sill, fitted sill and correlation length scale, with the practical correlation range defined as the distance at which the model reached 95% of its sill. Full mathematical details are provided in the Supplementary Information.

To visualise the spatial scale and amplitude of heterogeneity captured by the fitted model, Gaussian random field maps were generated from the fitted variogram parameters. These maps were used only as synthetic representations of the fitted spatial statistics and were not treated as experimental images. As a randomness control, pixel-shuffled maps were generated by randomly permuting relative-density values within the biological mask. Experimental maps were compared with shuffled controls using the positive autocorrelation area under the curve, expressed as a z-score relative to the shuffled distribution. Positive z-scores indicate stronger positive spatial autocorrelation than expected from random rearrangement of the same intensity values.

### Spontaneous Ca^2+^ release simulation

Ca^2+^ wave propagation was simulated on experimentally derived two-dimensional RyR/JPH2 maps. RyR positions were encoded as single-pixel objects with a fixed intensity value, while JPH2-positive junctional patches were encoded as non-zero, non-RyR pixels whose values represented the estimated local number of JPH2 labels. RyRs were classified as junctional only if they were located within a defined neighbourhood of a non-zero JPH2 patch. Non-junctional RyRs were excluded from the simulation. A binary JPH2 junctional mask, including retained RyR pixels, was generated for quality control and downstream analysis.

For each junctional RyR, the local JPH2 patch value was used to modulate RyR Ca^2+^ sensitivity. Specifically, RyR sensitivity was scaled inversely with the local JPH2 value, relative to the median JPH2 value across the junctional RyR population, with user-defined lower and upper bounds to avoid unrealistically large sensitivity differences. The mathematical formulation of this scaling is described in the Supplementary Information.

Ca^2+^dynamics were simulated on a two-dimensional grid with a pixel sampling of 10 nm/pixel. At each timestep, Ca^2+^ was updated according to an effective reaction-diffusion model incorporating cytosolic diffusion, first-order removal/buffering, and local RyR-mediated Ca^2+^ release. Resting cytosolic Ca^2+^ was set to 0.10 µM. Model parameters were chosen so that propagating cytosolic Ca^2+^ waves reached approximately 0.6-1.0 µM, while highly local RyR-centred nanodomains could transiently reach approximately 70 µM. The detailed reaction-diffusion equations and numerical implementation are provided in the Supplementary Information.

RyR gating was modelled stochastically. One RyR was triggered at the start of each simulation iteration, and subsequent RyR opening depended on the local Ca2+ concentration, a Ca^2+^ activation threshold, a Hill-type Ca^2+^ dependence, and the JPH2-dependent sensitivity scaling. Open durations were sampled from a bounded distribution, after which RyRs entered an inactive state for the remainder of the iteration. To initiate spatially directed propagation, the reference seed was selected from a junctional RyR cluster on the lower regions of the image, and repeated stochastic iterations were seeded within a 1 µm radius of this reference site.

For each condition, multiple independent iterations were simulated to capture stochastic variability in recruitment and wave spread. Quantitative outputs were saved as a 16-bit TIFF stacks representing absolute Ca^2+^ concentration over a fixed 0-70 µM range.

Longitudinal and transverse propagation were analysed from the saved 16-bit Ca^2+^ stacks. Because cells were not always aligned with the image axes, the cell longitudinal axis was manually specified as an angle measured clockwise from 12 o’clock. Line scans were extracted along the longitudinal and transverse axes, and iteration-wise kymograph collages were generated to compare stochastic wave behaviour. Propagation velocities were estimated from Ca^2+^ arrival times along each axis, with longitudinal and transverse speeds reported separately for each simulation iteration. A detailed description of the simulation steps are included in the Supplementary Information section.

### Statistical analysis

Unless otherwise stated, cluster-level measurements were first averaged per image (or per cell) to avoid pseudo-replication. Group comparisons between control and MCT were performed using two-sided Mann-Whitney U tests, where datasets or distributions did not satisfy the conditions for parametric statistical tests and sample sizes were modest. Data are presented as boxplots showing median, interquartile range, and full range excluding outliers. Linear relations between qIndex, puncta count, and stoichiometry were assessed using ordinary least squares regression, with confidence and prediction intervals estimated by bootstrap resampling where indicated. Exact p-values are reported in the corresponding figure panels.

## Ethics

Experiments were performed according to the UK Animals (Scientific Procedures) Act of 1986 and approval from the UK Home Office approval (licence number 70/8399) and the University of Leeds Ethics committee. All animals were subjected to Schedule 1 prior to the obtaining of heart tissues used in this study.

## Author contributions

R.K.: performed data curation, data analysis, software development, writing (original draft), review and editing; M.E.H.: performed all experiments; data acquisition, curation, funding acquisition, validation, writing, review, and editing; E.W.: project administration, resources, and review; I.J.: conceptualization, data curation, formal analysis, funding acquisition, methodology, project administration, resources, software, supervision, validation, visualization, writing, review, and editing.

## Acknowledgements

The authors acknowledge the NSW Health Cardiovascular Research Leadership Grant (H23/67588) awarded to I.J., the Wellcome Award (grant no. 207684/Z/17/Z) made to I.J., and the Leeds Anniversary Research Scholarship awarded to M.E.H. The authors extend their gratitude to Dr Thomas Sheard for assistance with experiments, and Prof Christian Soeller and Dr David Baddeley for advice on adapting the PyME software.

## References

1 Cheng, H., Lederer, W. J. & Cannell, M. B. Calcium sparks: elementary events underlying excitation-contraction coupling in heart muscle. Science 262, 740–744 (1993). 10.1126/science.8235594

2 Baddeley, D. et al. Optical single-channel resolution imaging of the ryanodine receptor distribution in rat cardiac myocytes. Proc Natl Acad Sci U S A 106, 22275–22280 (2009). 10.1073/pnas.0908971106

3 Hou, Y., Jayasinghe, I., Crossman, D. J., Baddeley, D. & Soeller, C. Nanoscale analysis of ryanodine receptor clusters in dyadic couplings of rat cardiac myocytes. J Mol Cell Cardiol 80, 45–55 (2015). 10.1016/j.yjmcc.2014.12.013

4 Asghari, P. et al. Cardiac ryanodine receptor distribution is dynamic and changed by auxiliary proteins and post-translational modification. Elife 9 (2020). 10.7554/eLife.51602

5 Franzini-Armstrong, C., Protasi, F. & Ramesh, V. Shape, size, and distribution of Ca(2+) release units and couplons in skeletal and cardiac muscles. Biophys J 77, 1528–1539 (1999). 10.1016/S0006-3495(99)77000-1

6 Pinali, C., Bennett, H., Davenport, J. B., Trafford, A. W. & Kitmitto, A. Three-dimensional reconstruction of cardiac sarcoplasmic reticulum reveals a continuous network linking transverse-tubules: this organization is perturbed in heart failure. Circ Res 113, 1219–1230 (2013). 10.1161/circresaha.113.301348

7 Jayasinghe, I. et al. True Molecular Scale Visualization of Variable Clustering Properties of Ryanodine Receptors. Cell Rep 22, 557–567 (2018). 10.1016/j.celrep.2017.12.045

8 Walker, M. A. et al. On the Adjacency Matrix of RyR2 Cluster Structures. PLoS Comput Biol 11, e1004521 (2015). 10.1371/journal.pcbi.1004521

9 Song, L. S. et al. Orphaned ryanodine receptors in the failing heart. Proc Natl Acad Sci U S A 103, 4305–4310 (2006). 10.1073/pnas.0509324103

10 Sun, X. H. et al. Molecular architecture of membranes involved in excitation-contraction coupling of cardiac muscle. J Cell Biol 129, 659–671 (1995). 10.1083/jcb.129.3.659

11 Marx, S. O. et al. PKA phosphorylation dissociates FKBP12.6 from the calcium release channel (ryanodine receptor): defective regulation in failing hearts. Cell 101, 365–376 (2000). 10.1016/s0092-8674(00)80847-8

12 Gyorke, I., Hester, N., Jones, L. R. & Gyorke, S. The role of calsequestrin, triadin, and junctin in conferring cardiac ryanodine receptor responsiveness to luminal calcium. Biophys J 86, 2121–2128 (2004). 10.1016/S0006-3495(04)74271-X

13 Takeshima, H., Komazaki, S., Nishi, M., Iino, M. & Kangawa, K. Junctophilins: a novel family of junctional membrane complex proteins. Mol Cell 6, 11–22 (2000). 10.1016/s1097-2765(00)00003-4

14 van Oort, R. J. et al. Disrupted junctional membrane complexes and hyperactive ryanodine receptors after acute junctophilin knockdown in mice. Circulation 123, 979–988 (2011). 10.1161/CIRCULATIONAHA.110.006437

15 Guo, A. et al. Overexpression of junctophilin-2 does not enhance baseline function but attenuates heart failure development after cardiac stress. Proc Natl Acad Sci U S A 111, 12240–12245 (2014). 10.1073/pnas.1412729111

16 Macquaide, N. et al. Ryanodine receptor cluster fragmentation and redistribution in persistent atrial fibrillation enhance calcium release. Cardiovasc Res 108, 387–398 (2015). 10.1093/cvr/cvv231

17 Sheard, T. M. D. et al. Three-dimensional visualization of the cardiac ryanodine receptor clusters and the molecular-scale fraying of dyads. Philos Trans R Soc Lond B Biol Sci 377, 20210316 (2022). 10.1098/rstb.2021.0316

18 Kolstad, T. R. et al. Ryanodine receptor dispersion disrupts Ca(2+) release in failing cardiac myocytes. Elife 7 (2018). 10.7554/eLife.39427

19 Benoist, D., Stones, R., Drinkhill, M., Bernus, O. & White, E. Arrhythmogenic substrate in hearts of rats with monocrotaline-induced pulmonary hypertension and right ventricular hypertrophy. Am J Physiol Heart Circ Physiol 300, H2230–2237 (2011). 10.1152/ajpheart.01226.2010

20 Xie, Y. P. et al. Sildenafil prevents and reverses transverse-tubule remodeling and Ca(2+) handling dysfunction in right ventricle failure induced by pulmonary artery hypertension. Hypertension 59, 355–362 (2012). 10.1161/hypertensionaha.111.180968

21 Medvedev, R. et al. Nanoscale Study of Calcium Handling Remodeling in Right Ventricular Cardiomyocytes Following Pulmonary Hypertension. Hypertension 77, 605–616 (2021). 10.1161/HYPERTENSIONAHA.120.14858

22 Hurley, M. E. et al. A correlative super-resolution protocol to visualise structural underpinnings of fast second-messenger signalling in primary cell types. Methods 193, 27–37 (2021). 10.1016/j.ymeth.2020.10.005

23 Barentine, A. E. S. et al. An integrated platform for high-throughput nanoscopy. Nat Biotechnol 41, 1549–1556 (2023). 10.1038/s41587-023-01702-1

24 Baddeley, D., Cannell, M. B. & Soeller, C. Visualization of localization microscopy data. Microsc Microanal 16, 64–72 (2010). 10.1017/S143192760999122X

25 Jungmann, R. et al. Quantitative super-resolution imaging with qPAINT. Nat Methods 13, 439–442 (2016). 10.1038/nmeth.3804

26 Sheard, T. M. D. et al. Three-Dimensional and Chemical Mapping of Intracellular Signaling Nanodomains in Health and Disease with Enhanced Expansion Microscopy. ACS Nano 13, 2143–2157 (2019). 10.1021/acsnano.8b08742

27 Holmes, M. et al. Increased SERCA2a sub-cellular heterogeneity in right-ventricular heart failure inhibits excitation-contraction coupling and modulates arrhythmogenic dynamics. Philos Trans R Soc Lond B Biol Sci 377, 20210317 (2022). 10.1098/rstb.2021.0317

