## Supplementary Information for "Structural and functional heterogeneity in cardiac RyR signalling nanodomains revealed with quantitative single molecule mapping toolkit"

* These authors contributed equally

### 1. Supplementary Methods

#### 1.1. Variogram-based analysis of spatial heterogeneity

Spatial heterogeneity in rendered RyR2 and JPH2 density maps was analysed using an approach adapted for cardiomyocytes by Holmes et al^1^. Rendered images were analysed separately for each channel. Before analysis, a shared mask was generated from the combined RyR2 and JPH2 density maps to exclude empty background regions. These images were then binned to pixel sampling of 100, 250, 500 and 1,000 nm/pixel and normalised within the mask to produce relative-density fields with a mean intensity of one.

Spatial autocovariance was calculated using a masked FFT-based approach to correct for finite image size and irregular mask geometry. The radially averaged autocorrelation profile, ρ(r), was converted to an empirical semi-variogram according to:

$$\gamma(r)=\sigma^{2}[1-\rho(r)]$$

where $\sigma^{2}$ is the variance of the normalised density field within the mask. Empirical semi-variograms were fitted with a Gaussian variogram model:

$$\gamma(r)=c_{0}+\sigma_{s}^{2}\left( 1 - e^{-r^{2}/(2\lambda^{2})} \right)$$

where $c_{0}$ represents the nugget, $\sigma_{s}^{2}$ the partial sill, $c_{0}+\sigma_{s}^{2}$ the fitted sill, and $\lambda$ the correlation length-scale. The practical correlation range, $\lambda_{95}$, was defined as the distance at which the fitted variogram reached 95% of its sill:

$$\lambda_{95}=\lambda\sqrt{-2\ln(0.05)}$$

Gaussian random field maps were generated from the fitted variogram parameters to visualise the spatial scale and amplitude of heterogeneity captured by the model. These maps were used only as synthetic representations of fitted spatial statistics and were not treated as experimental images. As a randomness control, pixel-shuffled maps were generated by randomly permuting relative-density values within the biological mask. Input renderings were compared with their shuffled controls using the positive autocorrelation area under the curve (AUC), expressed as a z-score relative to the shuffled distribution. Positive z-scores indicate stronger positive spatial autocorrelation than expected from random rearrangement of the same intensity values.

#### 1.2. Simulation of spontaneous and propagating Calcium release

##### 1.2.1. Input RyR/JPH2 maps

Two-dimensional maps of ryanodine receptor type 2 (RyR2) and junctophilin-2 (JPH2)-associated junctional domains were used as the spatial substrate for calcium release simulations. Input images were supplied as single-channel TIFF images with a pixel size of 5 nm. RyR2 positions were encoded as single-pixel objects with a fixed intensity value of 10000. JPH2-positive junctional patch areas were encoded as non-zero pixels with values distinct from the RyR2 marker value, where the pixel value represented the estimated number of JPH2 localisations within that patch.

RyR2 pixels were detected by exact matching to the defined RyR2 intensity value. JPH2 patch pixels were defined as all non-zero pixels that were not RyR2	pixels. A RyR2 was classified as junctional if at least one JPH2 patch pixel was present within a local neighbourhood around the RyR2 pixel. Unless otherwise	stated, this neighbourhood radius was one image pixel. RyR2s not associated with a JPH2-positive patch were excluded from all subsequent simulation and	analysis. A binary JPH2 junctional mask was also generated, including both the JPH2-positive patch area and the retained junctional RyR2 pixels.

##### 1.2.2. JPH2-dependent RyR sensitivity scaling

For each retained junctional RyR2, the local JPH2 patch value was calculated from the surrounding non-zero, non-RyR2 pixels. The local patch value was summarised using the median value of the neighbouring JPH2 patch pixels. RyR2 calcium sensitivity was then scaled inversely with local JPH2 patch value according to:

$$S_{i}=\mathrm{clip}\left[ \left( \frac{J_{\mathrm{ref}}}{J_{i}} \right)^{\gamma} , S_{\min} , S_{\max} \right]$$

where $S_{i}$is the calcium sensitivity scale for RyR2 $i$, $J_{i}$is the local JPH2 patch value, $J_{\mathrm{ref}}$is the median JPH2 value across retained junctional RyR2s, $\gamma$is the JPH2 sensitivity exponent, and $S_{\min}$and $S_{\max}$define lower and upper bounds on the scaling. In the final simulations, $\gamma=0.5$, $S_{\min}=0.5$, and $S_{\max}=2.0$. This scaling allowed RyR2 excitability to vary with the local nanoscale JPH2 environment while avoiding unrealistically large sensitivity differences.

##### 1.2.3. Simulation domain

The calcium simulation was performed on a two-dimensional Cartesian grid with 10 nm spacing. RyR2 coordinates from the 5 nm input image were mapped onto the simulation grid by nearest-neighbour coordinate conversion. The simulation domain was defined either from the full image or from a user-specified region of interest, with additional padding to avoid edge effects. Edges of the simulation grid were treated as absorbing boundaries through zero-padding during diffusion operations.

Each simulation iteration represented one stochastic release event initiated from a single triggered RyR2. Unless otherwise stated, simulations were run for 3000 ms with a timestep of 0.5 ms. Output frames were saved at 1 ms intervals.

##### 1.2.4. Seed selection

For simulations of propagating events, the reference trigger was selected from the lower-left region of the junctional RyR2 map. The binary JPH2/RyR2 junctional mask was segmented into connected components, and components within the leftmost fraction of the image were considered candidate seed clusters. The lower-most eligible cluster was selected, and the initial reference RyR2 seed was chosen from this cluster. For repeated stochastic iterations, seed RyR2s were selected within a 1 µm radius of the reference seed, allowing repeated initiation from the same local region while preserving trial-to-trial variability.

##### 1.2.5. Calcium reaction-diffusion model

The simulation represented cytosolic calcium concentration as an effective two-dimensional field. Calcium was modelled as the sum of resting calcium and release-evoked calcium:

$$C\left( x , y , t \right)=C_{\mathrm{rest}}+c\left( x , y , t \right)$$

where $C_{\mathrm{rest}}=0.10\text{ }\mu M$, corresponding to approximately 100 nM resting cytosolic calcium, and $c\left( x , y , t \right)$is the simulated calcium increase above rest.

At each timestep, the calcium field was updated by diffusion, decay, and local RyR2 release:

$$\frac{\partial c}{\partial t}=D_{\mathrm{eff}}\nabla^{2}c-\frac{c}{\tau_{\mathrm{decay}}}+R\left( x , y , t \right)$$

where $D_{\mathrm{eff}}$is the effective calcium diffusion coefficient, $\tau_{\mathrm{decay}}$is an effective calcium removal/buffering time constant, and $R\left( x , y , t \right)$represents calcium release from open RyR2s. In the final smooth-wavefront simulations, $D_{\mathrm{eff}}=0.006\text{ }\mu m^{2}\mathrm{ms}^{-1}$and $\tau_{\mathrm{decay}}=45\text{ }\mathrm{ms}$.

Diffusion was implemented numerically as a Gaussian convolution at each timestep, with:

$$\sigma_{\mathrm{diff}}=\frac{\sqrt{2D_{\mathrm{eff}}\Delta t}}{\Delta x}$$

where $\Delta t$is the simulation timestep and $\Delta x$is the simulation pixel size. The exponential decay term was implemented as multiplication by:

$$\exp\left( -\frac{\Delta t}{\tau_{\mathrm{decay}}} \right)$$

This formulation provides an effective cytosolic spread-and-clearance model rather than an explicit model of calcium buffers, pumps, or sarcoplasmic reticulum refilling.

##### 1.2.6. RyR calcium release

Each open RyR2 contributed a local calcium source to the simulation grid. Calcium release was added as a Gaussian-distributed source centred on the RyR2 position, with a source width of 120 nm. The peak release strength was set to 8.0 $\mu M\text{ }\mathrm{ms}^{-1}$per open RyR2 in the final smooth-wavefront simulations. Density-dependent RyR2 release amplification was disabled in the final simulations to avoid artificially reinforcing highly clustered regions and to favour smooth inter-cluster calcium propagation.

The simulation was calibrated so that the broad cytosolic calcium wave reached approximately 0.6–1.0 $\mu M$, while highly local RyR2-centred nanodomain values could transiently approach approximately 70 $\mu M$within the RyR2 pixel and immediately adjacent pixels.

##### 1.2.7. Stochastic RyR gating

Each RyR2 was modelled as a stochastic channel with discrete states: closed, open, and inactive. At the start of each iteration, the selected seed RyR2 was forced open at the trigger time. Other RyR2s opened stochastically according to a thresholded Hill-like Ca^2+^ recruitment rule, in which local Ca^2+^ above an activation threshold was raised to a Hill-like exponent, scaled by the local JPH2-dependent sensitivity factor, and capped at a maximum opening rate.

For each closed RyR2 $i$, the local calcium concentration $C_{i}$was sampled at the RyR2 grid position. The Ca^2+^ drive above threshold was defined as:

$${Ca}_{drive,i}=max({Ca}_{i}- {Ca}_{thr,i},0)$$

The opening rate was then calculated as:

$$k_{open, i}=\min\left[ k_{\max} , AS_{i}{Ca}_{drive,i}^{h} \right]$$

where $C_{i}$is the local Ca²⁺ concentration at RyR2 $i$, $C_{\mathrm{thr},i}$is the Ca²⁺ activation threshold, $Ca_{\mathrm{drive},i}$ is the Ca²⁺ concentration above threshold, $A$is the opening-rate coefficient, $S_{i}$is the JPH2-dependent sensitivity scale, $h$is the Hill-like exponent, and $k_{\max}$is the maximum allowed opening rate. The exponent $h$was applied to the Ca²⁺ drive above threshold, not to the total local Ca²⁺ concentration.

The probability of opening during a timestep was calculated as:

$$P_{open,i}=1-e^{-k_{open,i}\Delta t}$$

where is the simulation timestep. Channel opening was then determined by Monte Carlo sampling. In the final smooth-wavefront simulations, $A=0.0015$, $h=2.0$, $C_{\mathrm{thr}}=2.0 \mu M$, and $k_{\max}=0.025 ms^{-1}$. Open durations were sampled from a normal distribution with mean 22 ms and standard deviation 6 ms, bounded between 5 ms and 60 ms. After closure, RyR2s were placed into an inactive state and were not allowed to reopen during the same iteration.

##### 1.2.8. Rendering and image output

Quantitative calcium outputs were saved as 16-bit TIFF stacks representing absolute calcium concentration. Both raw and TIRF-like outputs were generated. The raw output preserved the simulation grid calcium values, while the TIRF-like output was generated by Gaussian spatial smoothing using a 200 nm point-spread approximation.

The quantitative 16-bit TIFF stacks used a fixed absolute scaling, with 0–70 $\mu M$ mapped across the 16-bit dynamic range. This preserved the full range from resting cytosolic calcium through RyR2-local nanodomain peaks. In addition, RGB TIFF stacks were generated using the same fixed linear calcium colour scale. No frame-by-frame intensity normalisation, percentile stretching, or colour-scale pinning was applied.

##### 1.2.9. Longitudinal and transverse wave propagation analysis

Propagation was analysed from the saved 16-bit raw calcium TIFF stacks. Because cardiomyocytes were not always aligned with the image axes, the cell longitudinal axis was specified manually as an angle measured clockwise from 12 o’clock. This angle defined the longitudinal unit vector, and the transverse unit vector was defined as the perpendicular direction.

For each simulation iteration, line scans were sampled through the calcium stack along the longitudinal and transverse axes. The line-scan centre was either user-defined or estimated from the calcium signal. A finite-width strip around each axis was averaged to reduce pixel-level noise. Line scans were saved as raw 16-bit kymograph TIFFs and as annotated RGB collages. In each collage, individual simulation iterations were arranged as separate row blocks to allow direct visual comparison of stochastic propagation patterns. Annotated outputs included calcium colour scale, spatial scale, and time scale indicators.

Wavefront propagation speeds were estimated from threshold-based arrival times along each line scan. For each spatial position, the arrival time was defined as the first time point at which the calcium signal crossed a defined fraction of the iteration-specific wave amplitude after baseline correction. Distance from the seed point was then regressed against arrival time separately on the positive and negative sides of each axis. The slope of this relationship provided the propagation velocity in $\mu m\text{ }s^{-1}$. Longitudinal and transverse speeds were reported separately for each iteration, together with fit quality metrics and the number of spatial points used for each fit.

1	Holmes, M. *et al.* Increased SERCA2a sub-cellular heterogeneity in right-ventricular heart failure inhibits excitation-contraction coupling and modulates arrhythmogenic dynamics. *Philos Trans R Soc Lond B Biol Sci* **377**, 20210317 (2022). <https://doi.org:10.1098/rstb.2021.0317>
